# Trends in outpatient antibiotic prescribing practice among US older adults, 2011-2015: an observational study

**DOI:** 10.1101/292243

**Authors:** Scott W. Olesen, Michael L. Barnett, Derek R. MacFadden, Marc Lipsitch, Yonatan H. Grad

## Abstract

**Objective:** To identify temporal trends in outpatient antibiotic use and antibiotic prescribing practice among older adults.

**Design:** Observational study using United States Medicare administrative claims during 2011-2015. Trends in antibiotic use were estimated using multivariable regression adjusting for beneficiaries’ demographic and clinical covariates.

**Setting:** Medicare.

**Participants:** 4.6 million Medicare beneficiaries from a nationwide, 20% sample of fee-for-service Medicare beneficiaries ≥65 years old.

**Main outcome measurements:** Overall rates of antibiotic prescription claims, rates of appropriate and inappropriate prescribing, rates for each of the most frequently prescribed antibiotics, and rates of antibiotic claims associated with specific diagnoses.

**Results:** Antibiotic claims fell from 1362.2 to 1361.6 claims per 1,000 beneficiaries per year during 2011-2015, an overall 0.2% decrease (95% CI 0.07-0.32). Inappropriate antibiotic claims fell from 552 to 533 claims per 1,000 beneficiaries, a 4.1% decrease (CI 3.9-4.3). Individual antibiotics had heterogeneous changes in use. For example, azithromycin claims per beneficiary decreased by 18.4% (CI 18.2-18.7) while levofloxacin claims increased by 28.1% (CI 27.5-28.6). Azithromycin use associated with each of the potentially appropriate and inappropriate respiratory diagnoses we considered decreased, while levofloxacin use associated with each of those diagnoses increased.

**Conclusion:** Among US Medicare beneficiaries, overall antibiotic use and inappropriate use declined modestly, but individual drugs experienced divergent changes in use. Trends in drug use across indications were stronger than trends in use for individual indications, suggesting that guidelines and concerns about antibiotic resistance were not major drivers of change in antibiotic use.

## Introduction

In the US, high rates of antibiotic prescribing pose a major challenge to public health.^1,2^ Clinically inappropriate antibiotic prescribing comprises a large fraction of overall use and contributes to increasingly broad antibiotic resistance.^3^ Despite guidelines and calls by federal agencies, professional medical societies, and other organizations to reduce antibiotic prescribing for inappropriate indications,^4^ prescribing patterns have changed little.^5–7^ Furthermore, data on antibiotic prescribing trends that can guide stewardship efforts remain sparse, particularly for key vulnerable populations, such as older adults.

Older adults are a particularly important population with respect to antibiotic overuse. They use approximately 50% more antibiotics per capita than younger adults^8^ and have the highest risk of poor outcomes from the adverse effects of antibiotics, including *Clostridium difficile* infection.^9^ After several decades of steady antibiotic use,^10–12^ during which use of individual drugs has varied,^11,13,14^ overall antibiotic use has begun to decline.^4,15–17^ Recent trends in antibiotic use among older adults, however, are unclear, perhaps having hit a peak in 2006 and a trough in 2014.^18,19^ Furthermore, there is limited data on trends for individual antibiotics and on use of individual antibiotics in association with specific indications. More definitive evidence on antibiotic use and its appropriateness in older adults is needed to guide stewardship interventions in this critical population.

To address this gap, we investigated recent trends in antibiotic prescribing among older adults, using administrative claims from the Medicare program, which provides healthcare insurance for most Americans over 65 years old, from 2011-2015. We focused specifically on trends in potentially inappropriate antibiotic use and heterogeneity in trends among individual antibiotics to identify potential targets for future stewardship interventions.

## Methods

### Study population

We studied a 20% sample of beneficiaries enrolled in Medicare outpatient medical insurance (Part B) and prescription coverage (Part D) for 2011 through 2015. For each data year, we included only individuals who were continuously enrolled in fee-for-service Medicare (i.e., no months in Medicare Advantage) for the entire year and who were at least 65 years old.

### Demographic and clinical variables

To characterize the differences in antibiotic use among older adults, we captured beneficiary sex, race/ethnicity, age, US Census region, eligibility for Medicaid (“dual eligibility”), and the presence of 20 chronic conditions (see Appendix).

### Classifying antibiotic claims

We examined original (i.e., not refill) outpatient prescription pharmacy claims for oral and injected antibiotics (Appendix Table 1) as defined using the Medicare Formulary file and aggregated by generic antibiotic formulation (Appendix Table 2). We excluded refills because we did not expect them to have associated prescriber encounters. We treated multiple claims from the same beneficiary on the same day for the same generic antibiotic as a single claim. To determine how individual antibiotics contributed to overall use and trends, we examined both overall antibiotic claims and claims for each of the 10 most frequently prescribed antibiotics. Medicare Part D claims data do not include information about antibiotics dispensed in inpatient facilities such as hospitals or skilled nursing facilities.

### Encounters and diagnoses

We linked antibiotic claims with outpatient prescriber encounters (Carrier and Outpatient files) and inpatient encounters (Inpatient and Skilled Nursing Facility files). An antibiotic claim was linked with an outpatient encounter if the prescription claim occurred on the day of the encounter or up to 7 days after. A claim was linked with an inpatient encounter if the claim was on the discharge date or up to 7 days after.

Diagnosis codes recorded for these encounters were grouped into 20 diagnostic categories (e.g., pneumonia) using a previously published classification scheme from a US Centers for Disease Control and Prevention workgroup, described in Fleming-Dutra *et al.*^2^ (We did not follow the exception listed in the “Bronchitis, bronchiolitis” category, “Excludes visits in which the 2nd or 3rd diagnosis was chronic bronchitis, emphysema, or COPD”, because this exception was specific to the coding format in their data source.^20^) We also followed that study’s organization of diagnostic categories into three antibiotic-appropriateness tiers: Tier 1 (“antibiotics almost always indicated”), Tier 2 (“antibiotics may be indicated”), and Tier 3 (“antibiotics not indicated”).

We classified antibiotic claims with an associated Tier 1 or Tier 2 diagnosis as appropriate. We classified claims with only Tier 3 diagnoses as inappropriate. We did not classify antibiotics without associated diagnoses as appropriate or inappropriate. This definition of appropriateness differs from the one used in by Fleming-Dutra *et al*. In that study, Tier 1 diagnoses were appropriate, most Tier 3 diagnoses were inappropriate, and appropriate antibiotic prescribing rates for the remaining diagnoses were determined using surveys of pathogen carriage or the minimum prescribing rate across the four US Census regions. We used a different definition of inappropriate use because pathogen carriage and geographical differences in prescribing rates themselves change with time and are thus confounded with temporal trends in appropriate use.

In October 2015, the US healthcare system, including Medicare records, transitioned from ICD-9 to ICD-10 diagnosis codes, but the scheme in Fleming-Dutra *et al.* uses ICD-9 codes. To avoid bias from coding changes, we did not link antibiotic claims in the fourth quarter of 2015 with diagnoses.

### Outcomes

To assess trends in antibiotic use, the outcome of interest was the number of antibiotic claims per 1,000 beneficiaries in each year. We considered total antibiotic claims, appropriate claims, inappropriate claims, claims without associated diagnoses, and claims for each of the 10 frequently prescribed antibiotics. Because claims in the fourth quarter of 2015 were not linked to diagnoses, rates of appropriate claims, inappropriate claims, and claims without associated diagnoses were projected from the first three quarters to the full year (by dividing by 0.75) for 2015 only.

To assess the contribution of individual diagnoses to these trends, the outcome of interest was the number of claims for a particular drug linked with a particular diagnosis (e.g., number of azithromycin claims associated with a pneumonia diagnosis) per 1,000 beneficiaries in a year. We considered the 13 diagnosis categories with the most associated antibiotic claims, grouped those diagnosis categories into 4 infection sites, and considered the 3 antibiotics contributing the most claims for each infection site (see Appendix).

### Statistical analyses

To assess trends in the study population characteristics, we performed Poisson regressions (number of beneficiaries, mean age, mean number of chronic conditions) or log-binomial regressions (proportion female, white, dual eligible, and in each Census region) predicting the population characteristic from study year. Adjusted trends in antibiotic use (i.e., claims per beneficiary per year) were assessed using Poisson regression. The main covariate of interest was a linear term for year. We adjusted for beneficiary age, sex (male or female), race (white, black, Hispanic, other), Census region, dual eligibility, and number of chronic conditions. For regressions involving appropriate claims, inappropriate claims, and claims without diagnoses, the period of exposure for each of 2011-2014 was 1 year and for 2015 was 9 months (i.e., excluding October-December during which ICD-10 was in use), and an offset term was included. When reporting a trend in use, the coefficient for year was projected to the full 2011-2015 span for all regressions. Regressions were performed using PROC GENMOD in SAS (version 9.4). Clustered standard error estimators accounting for correlations between multiple measurements from the same beneficiaries^21^ yielded similar confidence intervals.

## Results

Our study sample included 4.6 million unique beneficiaries with 19.8 million antibiotic claims from 2011-2015 (Table 1). Population characteristics changed by less than 10% over the study period except for the number of beneficiaries and proportion of beneficiaries eligible for Medicaid (Appendix Table 3).

**Table 1.** Study population characteristics.

|  | Year |  |  |  |  |
| --- | --- | --- | --- | --- | --- |
|  | 2011 | 2012 | 2013 | 2014 | 2015 |
| No. beneficiaries | 2,479,672 | 2,622,891 | 3,039,105 | 3,185,933 | 3,474,641 |
| Mean age (s.d.) | 75.5 (7.7) | 75.4 (7.7) | 75.2 (7.7) | 75.0 (7.7) | 75.2 (7.8) |
| Mean no. of chronic conditions (s.d.) | 2.70 (1.83) | 2.69 (1.83) | 2.69 (1.82) | 2.67 (1.82) | 2.72 (1.88) |
| % female | 63.1 | 62.4 | 61.3 | 61.3 | 62.6 |
| % white | 80.3 | 80.5 | 81.3 | 81.9 | 82.6 |
| % dual eligible | 26.4 | 25.0 | 21.3 | 19.5 | 17.9 |
| Region (%) |  |  |  |  |  |
| South | 38.7 | 38.3 | 37.9 | 38.1 | 38.1 |
| Midwest | 24.3 | 23.8 | 24.3 | 23.9 | 23.5 |
| West | 18.5 | 18.6 | 18.2 | 18.2 | 18.3 |
| Northeast | 18.5 | 19.3 | 19.6 | 19.8 | 20.2 |

Use of all antibiotics fell from 1,362.2 claims per 1,000 beneficiaries in 2011 to 1,361.6 claims per 1,000 beneficiaries in 2015, an overall 0.20% decrease (95% CI 0.07-0.32, *p* = 0.002; Figures 1a, 2). Trends in use varied by age, race, and region, and use actually increased among beneficiaries aged 76 to 85, female beneficiaries, white beneficiaries, and beneficiaries in the South and Northeast (*p* < 0.001 for all trends; Appendix Table 4).

**Figure 1:**
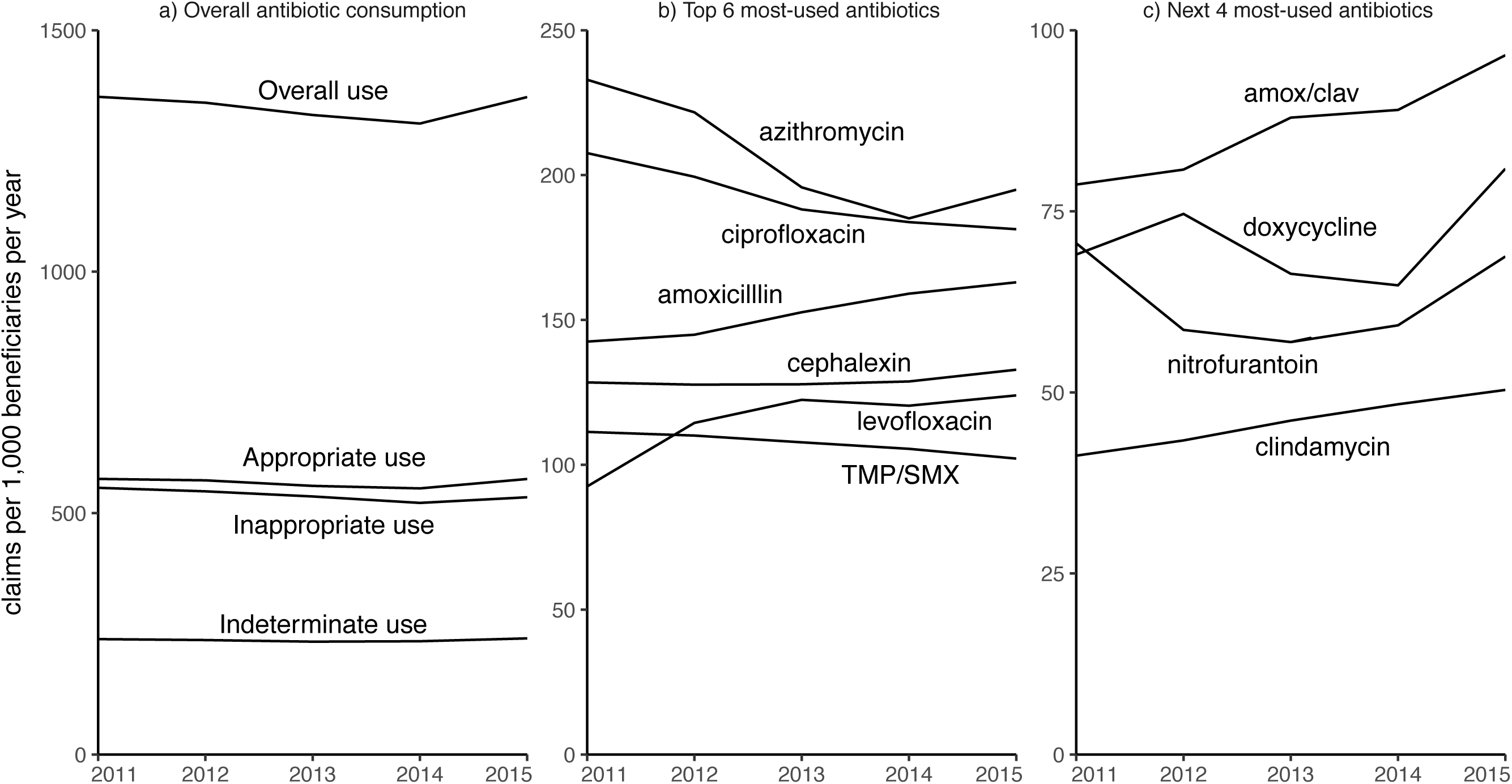
Rates of antibiotic prescribing, 2011-2015. Lines indicate claims per 1,000 beneficiaries per year for all antibiotic claims, inappropriate claims, appropriate claims, and claims without associated diagnoses (“Indeterminate use”; 1a); and for claims for each of the 10 most-claimed antibiotics (1b and 1c). Inappropriate, appropriate, and indeterminate use in 2015 are projected from the first three quarters to a full year. TMP/SMX: trimethoprim/sulfamethoxazole. amox/clav: amoxicillin/clavulanate.

Overall inappropriate prescribing fell from 552 claims per 1,000 beneficiaries in 2011 to 533 claims per 1,000 beneficiaries in 2015, a 4.1% decrease (95% CI 3.9-4.3, *p* < 0.001; Figure 2), while overall appropriate prescribing was at 570 claims per 1,000 beneficiaries in both 2011 and 2015 (0.8% increase, 95% CI 0.6-1.0, *p* < 0.001; Figure 2). The proportion of claims that were inappropriate declined from 40.5% in 2011 to 39.6% in 2015.

**Figure 2:**
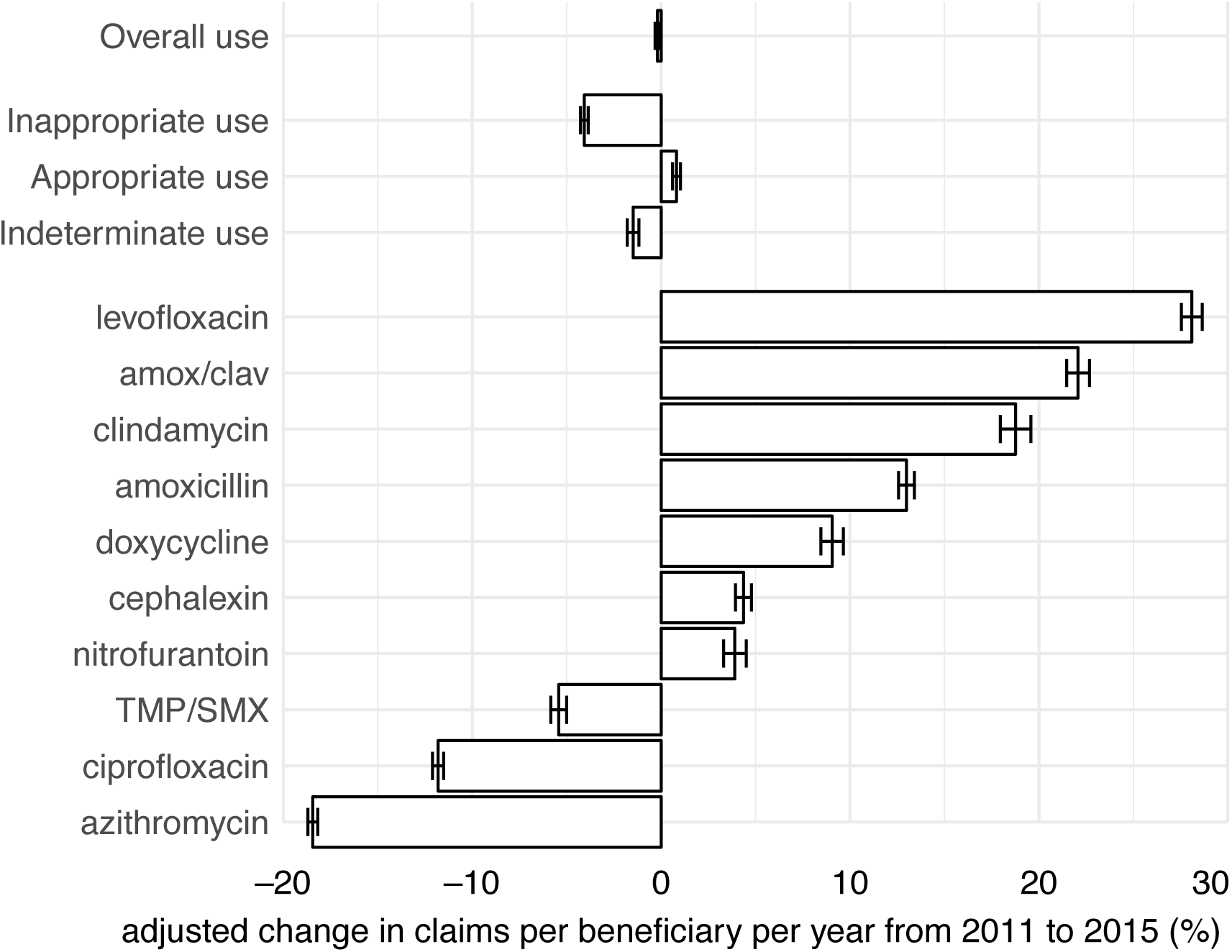
Adjusted trends in antibiotic use, 2011-2015. Bars indicate relative changes in claims per beneficiary per year for all antibiotic claims; for inappropriate claims, appropriate claims, and claims without associated diagnoses (“Indeterminate use”); and for claims for each of the 10 most-claimed antibiotics. Relative changes were determined by Poisson regression on claims per beneficiary per year adjusted for age, sex, race, census region, dual eligibility, and number of chronic conditions. Error bars show 95% confidence intervals. *p* < 0.001 for all trends (Wald test).amox/clav: amoxicillin/clavulanate. TMP/SMX: trimethoprim/sulfamethoxazole.

The 10 most frequently prescribed antibiotics, which accounted for 86.2% of all antibiotic claims, showed heterogeneous trends (Figures 1b, 1c, 2). Use declined for azithromycin, ciprofloxacin, and trimethoprim/sulfamethoxazole, but use rose for the 7 other drugs. The most marked changes were in azithromycin (decline of 18.4%, 95% CI 18.2-18.7, *p* < 0.001) and levofloxacin (increase of 28.1%, 95% CI 27.5-28.6, *p* < 0.001).

For levofloxacin, amoxicillin/clavulanate, and azithromycin, trends in use were similar across antibiotic-appropriate and antibiotic-inappropriate respiratory diagnoses. Use of levofloxacin, which increased overall, also increased in association with each of the appropriate and inappropriate respiratory diagnostic categories we considered: pneumonia, sinusitis, viral upper respiratory tract infections, bronchitis, asthma and allergy, and other respiratory conditions (Table 2). Use of amoxicillin/clavulanate, which increased overall, also increased in association with each of these respiratory diagnoses, except pneumonia, for which we did not detect a trend. In contrast, use of azithromycin, which decreased overall, decreased in association with each of these appropriate and inappropriate diagnostic categories.

**Table 2.** Trends in use of azithromycin, levofloxacin, and amoxicillin/clavulanate for respiratory conditions

|  |  | Claims per 1,000 beneficiaries per year<sup>a</sup> |  |  |  |  |  |  |  |  |
| --- | --- | --- | --- | --- | --- | --- | --- | --- | --- | --- |
|  |  | Azithromycin |  |  | Levofloxacin |  |  | Amoxicillin/clavulanate |  |  |
| Diagnosis | Antibiotics appropriate?<sup>b</sup> | 2011 | 2015<sup>c</sup> | Relative change<sup>d</sup> (%) | 2011 | 2015<sup>c</sup> | Relative change<sup>d</sup> (%) | 2011 | 2015<sup>c</sup> | Relative change<sup>d</sup> (%) |
| Pneumonia<sup>e</sup> | Yes | 17.4 | 14.4 | -18.3 ± 1.1 | 28.6 | 37.0 | 25.2 ± 1.1 | 8.8 | 8.7 | -0.1 ± 1.7 (n.s.) |
| Sinusitis | Potentially | 29.6 | 24.7 | -21.1 ± 0.8 | 6.9 | 12.5 | 59.0 ± 2.4 | 15.3 | 23.1 | 49.9 ± 1.8 |
| VURI<sup>f</sup> | No | 60.1 | 56.0 | -9.1 ± 0.6 | 19.2 | 31.1 | 53.1 ± 1.5 | 10.8 | 15.6 | 43.5 ± 2.0 |
| Acute bronchitis<sup>g</sup> | No | 56.1 | 46.4 | -19.9 ± 0.6 | 16.1 | 25.3 | 42.8 ± 1.5 | 8.6 | 10.3 | 18.2 ± 1.9 |
| Other respiratory<sup>h</sup> | No | 61.1 | 50.8 | -16.9 ± 0.6 | 55.7 | 69.5 | 22.6 ± 0.8 | 23.4 | 24.9 | 7.4 ± 1.1 |
| Asthma & allergy<sup>i</sup> | No | 21.6 | 19.8 | -10.6 ± 1.0 | 8.4 | 12.7 | 41.7 ± 2.1 | 5.9 | 7.9 | 34.4 ± 2.6 |

a: Includes claims for each antibiotic linked to the given diagnosis. A single claim may be linked to multiple diagnoses, so the sum of the “2011” and “2015” columns exceeds the number of claims associated with any listed diagnosis (see Appendix).
b: As determined by the CDC working group<sup>2</sup> (see Methods)
c: Claims in the fourth quarter of 2015 were not linked to diagnoses. These values are projected from the first three quarters to a full-year period (see Methods).
d: Adjusted value for 2011-2015 change from Poisson regression on claims per beneficiary per year adjusted for age, sex, race, census region, dual eligibility, and number of chronic conditions. Plus-minus contains 95% confidence interval. $p < 0.001$ (Wald $\chi^2$ test) for all trends except pneumonia and amoxicillin/clavulanate ( $p = 0.89$ ).
e: Includes *Streptococcus pneumoniae* pneumonia, other bacterial pneumonia, pneumonia due to other specified organism, pneumonia in infectious diseases classified elsewhere, bronchopneumonia with organism unspecified, pneumonia with organism unspecified
f: Includes acute nasopharyngitis, acute laryngitis and tracheitis, acute upper respiratory infections of multiple or unspecified sites, and cough
g: Includes bronchitis not specified as acute or chronic, acute bronchitis and bronchiolitis
h: Includes chronic bronchitis, dyspnea, stridor, hemoptysis, and abnormal sputum
i: Includes allergic rhinitis and unspecified allergy

For example, use of azithromycin following pneumonia diagnoses, for which antibiotics are considered appropriate, fell from 17.4 claims per 1,000 beneficiaries in 2011 to 14.4 claims per 1,000 beneficiaries in 2015, an 18.3% decrease (95% CI 17.2-19.3, *p* < 0.001), and use of azithromycin following viral upper respiratory tract infection, for which antibiotics are considered inappropriate, fell from 60.1 to 56.0 claims per 1,000 beneficiaries per year, a 9.1% decrease (95% CI 8.5-9.7, *p* < 0.001). In contrast, use of levofloxacin for pneumonia rose from 28.6 to 37.0 claims per 1,000 beneficiaries, a 25.2% increase (95% CI 24.1-26.2, *p* < 0.001), and use of levofloxacin for viral upper respiratory tract infection rose from 19.2 to 31.1 claims per 1,000 beneficiaries, a 53.1% increase (95% CI 51.7-54.6, *p* < 0.001).

Azithromycin use in association with any of these categories declined from 145.7 to 123.4 claims per 1,000 beneficiaries per year (a decline of 22.3), while levofloxacin use increased from 53.3 to 77.0 claims per 1,000 beneficiaries per year (an increase of 23.7). Thus, the decline in azithromycin use for respiratory conditions was exceeded by the increases in levofloxacin use and was not associated with a net decrease in antibiotic prescribing for respiratory conditions.

Prescribing practice patterns for gastrointestinal infections, for which antibiotics are potentially appropriate, and other gastrointestinal conditions, for which antibiotics are inappropriate, displayed a similar pattern of use across antibiotic-appropriate and antibiotic-inappropriate diagnoses (Appendix Table 5). Use of ciprofloxacin, which fell overall, also fell in association with both infections and other conditions, while use of levofloxacin, which rose overall, also rose in association with both infections and other conditions. Use of metronidazole, which decreased overall (5.1%, 95% CI 4.3-5.8, *p* < 0.001), fell in association with infections, but we did not detect a trend in use of metronidazole associated with other gastrointestinal conditions. Prescribing practice for genitourinary and skin/cutaneous/mucosal conditions also largely displayed the pattern of increases or decreases by antibiotic across indications, regardless of appropriateness (Appendix Table 6, Appendix Table 7).

## Discussion

In this analysis of a national sample of Medicare beneficiaries from 2011 to 2015, we found that overall antibiotic use declined a modest 0.2% and overall inappropriate prescribing declined 4.2%. There was increased use of levofloxacin over azithromycin for all respiratory conditions during this period, whether antibiotics were appropriate for that indication or not. Prescribing practice for other conditions displayed similar patterns, suggesting that changes in antibiotic prescribing practice may be due to shifting preferences for antibiotics across indications rather than targeted reductions in use of particular antibiotics for particular indications. Furthermore, the proportion of antibiotic use we observed as inappropriate (approximately 40%) was substantially higher than previously reported^2^ (18% for ≥65 years old), indicating that inappropriate antibiotic prescribing among Medicare beneficiaries may be much higher than expected.

There are at least four explanations for the trends we observed in the use of individual antibiotics. Our results are inconsistent with the first two but consistent with the second two. First, trends could represent altered prescribing practice in response to changed prescribing guidelines. For example, in 2007 the Infectious Diseases Society of America (IDSA) and the American Thoracic Society issued a recommendation that a respiratory fluoroquinolone be used instead of a macrolide for treating community-acquired pneumonia in certain patients.^22^ If this recommendation were still in the process of being implemented during 2011-2015, it could help explain why azithromycin use for pneumonia decreased 18% while use of levofloxacin for pneumonia increased 25% (Table 3). Second, the trends could reflect concerns about antibiotic resistance, such as rising macrolide resistance among community-acquired pneumonia,^23^ which could also explain decreased use of azithromycin for pneumonia. Third, the trends could reflect safety concerns about particular antibiotics. For example, in 2013 the US Food and Drug Administration warned prescribers that azithromycin might increase the risk of cardiovascular death,^24^ which may have contributed to the decreased popularity of azithromycin across diagnoses. Fourth, these trends might represent market factors, such as pricing, availability, and advertising, as suggested in a previous study^12^ that found that broad-spectrum antibiotics became more used for both antibiotic-appropriate and antibiotic-inappropriate respiratory indications during 1995-2002.

The first two of these explanations, guidelines and concerns about antibiotic resistance, should be specific to particular antibiotics and diagnoses, while the latter two, safety and market factors, should apply to antibiotics across diagnoses. In this study, we found that use of azithromycin for all respiratory conditions fell, while use of levofloxacin for those conditions rose. This pattern is inconsistent with diagnosis-specific explanations, suggesting that the observed trends in drug use are driven more by safety concerns or market factors than by altered guidelines or by concerns about antibiotic resistance. Furthermore, the slow decline in overall antibiotic use we observed, coupled with the fast trends in use of individual drugs, suggests it is easier to substitute one antibiotic for another than to reduce overall antibiotic use.^25,26^

Our study has several limitations. First, the fee-for-service service beneficiaries for which we have more complete prescription and encounter data may not be representative of the entire US older adult population.^27^ Second, pharmacy claims data do not contain information about the condition the drug is intended to treat, so our matching of encounters and prescriptions may have measurement bias or be affected by trends in coding practice.^28,29^ In addition, we were not able to classify approximately 18% of antibiotic claims as appropriate or inappropriate because they had no associated diagnoses, similar to a previous study^30^ that found that 15% of Medicare antibiotic claims had no associated encounter. Third, the 5-year period of our study prevents assessment of longer term trends. As a final caveat, we note that the difference between the inappropriate prescribing proportion in this study (40%) and in a previous report^2^ (18% among Americans at least 65 years old) is likely due to the different data sources and methods of linking antibiotic claims with diagnoses in the two studies. In the NAMCS/NHAMCS surveys^20^ used by Fleming-Dutra *et al*.,^2^ each antibiotic prescription is definitively associated with its motivating diagnosis. Furthermore, the exact definitions of inappropriate use differed between this study and Fleming-Dutra *et al*. (see Methods). The differences between our study and Fleming-Dutra *et al*. demonstrate the challenges in quantifying inappropriate antibiotic use and the need for definitions of appropriate antibiotic use that can be applied across data sources.

In conclusion, we find that overall antibiotic use and inappropriate antibiotic prescribing decreased modestly in a national population of older adults from 2011 to 2015. For drugs used to treat respiratory infections, most changes in antibiotic use were due to increased use of one antibiotic over another across indications, not clinically-oriented changes in use. Changes in antibiotic prescribing practice reflected shifting use between antibiotics rather than declining inappropriate prescribing across antibiotics. Thus, despite decades of effort, appropriateness of nationwide antibiotic use is improving at most incrementally. Bridging the large gap between our goals and past performance will require strong, national policy changes and innovative approaches to stewardship.

## Statements and permissions

### Details of the contributors

SWO, MLB and YHG conceived of and designed the work. SWO analyzed the data. SWO and YHG drafted the work. All authors revised the work critically and approved the final manuscript.

### Ethical approval

This study was deemed exempt from review by the institutional review board at the Harvard T. H. Chan School of Public Health, Boston, Massachusetts, USA.

### Data sharing

All data are available from the Centers for Medicare and Medicaid Services.

## References

1. Shapiro DJ, Hicks LA, Pavia AT, Hersh AL. Antibiotic prescribing for adults in ambulatory care in the USA, 2007–09. J Antimicrob Chemother. 2014 Jan 1;69(1):234–40.

2. Fleming-Dutra KE, Hersh AL, Shapiro DJ, Bartoces M, Enns EA, File TM, et al. Prevalence of Inappropriate Antibiotic Prescriptions Among US Ambulatory Care Visits, 2010-2011. JAMA. 2016 May 3;315(17):1864–73.

3. Gonzales R, Malone DC, Maselli JH, Sande MA. Excessive Antibiotic Use for Acute Respiratory Infections in the United States. Clin Infect Dis. 2001 Sep 15;33(6):757–62.

4. Centers for Disease Control and Prevention. Antibiotic Prescribing and Use in the US [Internet]. 2017 [cited 2018 Mar 19]. Available from: https://www.cdc.gov/antibioticuse/stewardship-report/index.html

5. Barnett ML, Linder JA. Antibiotic Prescribing to Adults With Sore Throat in the United States, 1997-2010. JAMA Intern Med. 2014 Jan 1;174(1):138–40.

6. Barnett ML, Linder JA. Antibiotic Prescribing for Adults With Acute Bronchitis in the United States, 1996-2010. JAMA. 2014 May 21;311(19):2020–2.

7. Schmidt ML, Spencer MD, Davidson LE. Patient, Provider, and Practice Characteristics Associated with Inappropriate Antimicrobial Prescribing in Ambulatory Practices. Infect Control Amp Hosp Epidemiol. 2018 Mar;391(3):307–15.

8. Hicks LA, Bartoces MG, Roberts RM, Suda KJ, Hunkler RJ, Taylor TH, et al. US outpatient antibiotic prescribing variation according to geography, patient population, and provider specialty in 2011. Clin Infect Dis Off Publ Infect Dis Soc Am. 2015 May 1;60(9):1308–16.

9. Lessa FC, Mu Y, Bamberg WM, Beldavs ZG, Dumyati GK, Dunn JR, et al. Burden of Clostridium difficile infection in the United States. N Engl J Med. 2015 Feb 26;372:825–34.

10. McCaig LF, Hughes JM. Trends in Antimicrobial Drug Prescribing Among Office-Based Physicians in the United States. JAMA. 1995 Jan 18;273(3):214–9.

11. McCaig LF, Besser RE, Hughes JM. Antimicrobial-Drug Prescription in Ambulatory Care Settings, United States, 1992–2000. Emerg Infect Dis [Internet]. 2003 [cited 2018 Mar 19];9(4). Available from: https://wwwnc.cdc.gov/eid/article/9/4/02-0268_article

12. Roumie CL, Halasa NB, Grijalva CG, Edwards KM, Zhu Y, Dittus RS, et al. Trends in antibiotic prescribing for adults in the United States—1995 to 2002. J Gen Intern Med. 2005 Aug 1;20(8):697.

13. Steinman MA, Gonzales R, Linder JA, Landefeld CS. Changing Use of Antibiotics in Community-Based Outpatient Practice, 1991-1999. Ann Intern Med. 2003 Apr 1;138(7):525.

14. Grijalva CG, Nuorti JP, Griffin MR. Antibiotic Prescription Rates for Acute Respiratory Tract Infections in US Ambulatory Settings. JAMA. 2009 Aug 19;302(7):758–66.

15. Suda KJ, Hicks LA, Roberts RM, Hunkler RJ, Matusiak LM, Schumock GT. Antibiotic Expenditures by Medication, Class, and Healthcare Setting in the United States, 2010–2015. Clin Infect Dis. 2018 Jan 6;66(2):185–90.

16. Van Boeckel TP, Gandra S, Ashok A, Caudron Q, Grenfell BT, Levin SA, et al. Global antibiotic consumption 2000 to 2010: an analysis of national pharmaceutical sales data. Lancet Infect Dis. 2014 Aug 1;14(8):742–50.

17. Blue Cross Blue Shield. Antibiotic prescription fill rates declining in the US [Internet]. 2017 [cited 2018 Mar 19]. Available from: https://www.bcbs.com/the-health-ofamerica/reports/antibiotic-prescription-rates-declining-in-the-US

18. Lee GC, Reveles KR, Attridge RT, Lawson KA, Mansi IA, Lewis JS, et al. Outpatient antibiotic prescribing in the United States: 2000 to 2010. BMC Med. 2014 Jun 11;12:96.

19. Durkin MJ, Jafarzadeh SR, Hsueh K, Sallah YH, Munshi KD, Henderson RR, et al. Outpatient Antibiotic Prescription Trends in the United States: A National Cohort Study. Infect Control Amp Hosp Epidemiol. 2018 Feb;1–6.

20. Centers for Disease Control and Prevention. Ambulatory Health Care Data [Internet]. [cited 2018 Mar 19]. Available from: https://www.cdc.gov/nchs/ahcd/index.htm

21. MacKinnon JG, White H. Some heteroskedasticity-consistent covariance matrix estimators with improved finite sample properties. J Econom. 1985 Sep 1;29(3):305–25.

22. Mandell LA, Wunderink RG, Anzueto A, Bartlett JG, Campbell GD, Dean NC, et al. Infectious Diseases Society of America/American Thoracic Society Consensus Guidelines on the Management of Community-Acquired Pneumonia in Adults. Clin Infect Dis. 2007 Mar 1;44(Supplement_2):S27–72.

23. Lynch JP, Martinez FJ. Clinical relevance of macrolide-resistant Streptococcus pneumoniae for community-acquired pneumonia. Clin Infect Dis. 2002 Mar 1;34 Suppl 1:S27–46.

24. Ray WA, Murray KT, Hall K, Arbogast PG, Stein CM. Azithromycin and the Risk of Cardiovascular Death. N Engl J Med. 2012 May 17;366(20):1881–90.

25. Burke JP. Antibiotic Resistance—Squeezing the Balloon? JAMA. 1998 Oct 14;280(14):1270–1.

26. Seppälä H, Klaukka T, Vuopio-Varkila J, Muotiala A, Helenius H, Lager K, et al. The Effect of Changes in the Consumption of Macrolide Antibiotics on Erythromycin Resistance in Group A Streptococci in Finland. N Engl J Med. 1997 Aug 14;337(7):441–6.

27. Riley GF, Levy JM, Montgomery MA. Adverse Selection In The Medicare Prescription Drug Program. Health Aff (Millwood). 2009 Nov 1;28(6):1826–37.

28. Department of Health and Human Services. Coding trends of Medicare evaluation and management services. 2012. Report No.: OEI-04-10-00180.

29. Pichichero ME. Dynamics of Antibiotic Prescribing for Children. JAMA. 2002 Jun 19;287(23):3133–5.

30. Riedle BN, Polgreen LA, Cavanaugh JE, Schroeder MC, Polgreen PM. Phantom Prescribing: Examining the Frequency of Antimicrobial Prescriptions Without a Patient Visit. Infect Control Amp Hosp Epidemiol. 2017 Mar;38(3):273–80.

